# Encoding social preference by interhemispheric neurons in the Insula

**DOI:** 10.1101/2022.12.15.520538

**Authors:** Christelle Glangetas, Elodie Ladevèze, Adriane Guillaumin, Manon Gauthier, Evelyne Doudnikoff, Erwan Bézard, Anne Taupignon, Jérôme Baufreton, François Georges

## Abstract

The Insula is a multisensory relay that participates in socio-emotional processing through multiple projections to sensory, cognitive, emotional, and motivational regions. Interestingly, the Insula interhemispheric projection to the contralateral Insula is a strong but understudied projection. Using cutting-edge neuroanatomy, *ex vivo* and *in vivo* electrophysiology associated with specific circuit manipulation, we unraveled the nature and role of Insula interhemispheric communication in social and anxiety processing in mice. In this study, we 1) characterized the anatomical and molecular profile of the interhemispheric neurons of the Insula, 2) highlighted that stimulation of this neuronal subpopulation triggers excitation in the Insula interhemispheric circuit 3) uncovered their engagement in social processing. In conclusion, this study demonstrates that interhemispheric neurons of the Insula constitute a unique class of Insula neurons and proposes new meaningful insights into the neuronal mechanisms underlying social behavior.

## Introduction

The Insular Cortex is classically described as an integrator of multimodal sensory signals coming from external cues (the environment) and internal cues (the body changes). For example, Insula responds to auditory or tactile cues^1^ and to cardiac interoceptive signals^2^. Interacting with novel individuals is an experience that leads to the integration of signals from both interoceptive and exteroceptive sources. Recently, it has been shown that some Insula cells respond to social interaction^3^. Interestingly these “social-on” cells solely represent a subset of Insula neurons that remained unexplored. In physiological situations, Insula neurons are engaged in social interaction and notably in social affective behaviors^4–9^. For example, it has been highlighted that Insula neurons projecting to the nucleus accumbens core regulate the social approach to stressed juvenile rats^8^. Autism spectrum Disorders (ASD) and Anxiety Disorders are pathologies with sensory integration defects that have been associated with dysfunction of the Insula^1,1,10–13^. An Insula maturation deficit was detected in a mouse model of ASD which is notably characterized by social interaction deficits, leading to an alteration in the integration of sensory information within the Insula^1^. Moreover, clinical studies show an Insula overactivation in anxious patients^10,14^. Altogether, these studies suggest that Insula is well-positioned to integrate and participate in regulating of socio-emotional processing. Indeed, the Insula shares multiple projections with sensory and interoceptive regions (sensory cortex, thalamus, olfactory bulb), with cognitive regions (medial prefrontal cortex, orbitofrontal cortex), emotional territories (amygdala, bed nucleus of the stria terminalis), and motivation-associated structures (ventral tegmental area, nucleus accumbens)^15^.

A strong but understudied projection is the Insula interhemispheric projection to the contralateral Insula^16^. As alteration in interhemispheric communication is associated with a social deficit^17–20^, we postulated that Insula interhemispheric communication is essential to develop adaptive reactions when facing novel social cues or threatening situations.

Recent evidence points toward a crucial role of cortical interhemispheric communication in complex cognitive and emotional processing. For example, individuals with high anxiety levels present an altered interhemispheric communication in reaction to the presentation of emotional images ^21^. Across mammalian evolution, cortical interhemispheric communication occurs notably through the corpus callosum^22^. Alterations in callosal fiber integrity have been observed in several pathological conditions as in patients with strokes, multiple sclerosis, schizophrenia, or ASD^23,24^. Despite recent advances in Insula participation in social behavior, the anatomical and molecular profile and the role of the Insula interhemispheric circuit in this socio-emotional processing remained poorly understood.

We hypothesized that Insula interhemispheric communication is essential to regulate social interactions and anxiety phenotype, and alteration in this communication would lead to social impairments and maladaptive anxiety behavior. To define the critical, yet unknown role of the Insula interhemispheric circuit, we used a combination of innovative neurotechniques, *in vivo* electrophysiology, and behavioral assays coupled with selective genetic neuron ablation and circuit manipulation in mice. This study was developed around 3 specific objectives: **i)** Anatomical and molecular characterization of Insula interhemispheric neurons, **ii)** Synaptic and circuit properties of Insula interhemispheric communication **iii)** Role of Insula interhemispheric communication in social interaction and anxiety-related behaviors in mice.

## Results

### Interhemispheric Insula neurons represent a unique subpopulation of the Insula

We first confirmed that Insula project to multiple brain regions by using an anterograde monosynaptic viral approach (Supp. Fig 1a). Interestingly, we observed a strong bilateral innervation to the dorsolateral part of the bed nucleus of the stria terminalis (dlBNST), the Central Amygdala (CeA), and contralateral labeling to the Insula (Supp. Fig 1b-f). To identify the projection targets of this Insula to-Insula circuit, we first mapped Insula interhemispheric neuron outputs, by injecting a retrograde monosynaptic virus (rAAV2-retro-CAG-Cre) in the contralateral Insula coupled with an anterograde monosynaptic virus (AAV2-DIO-eif1a-eYFP) injection in the ipsilateral Insula (Fig 1a). We showed that Insula interhemispheric neurons also projected massively to the dlBNST and the CeA (Fig 1b-g). In addition, we targeted the same interhemispheric Insula neuron population by injecting the retrograde monosynaptic virus in the CeA and the anterograde monosynaptic virus in the ipsilateral Insula. We confirmed that CeA-projecting Insula neurons also innervate both the dlBNST and the contralateral Insula (Supp Fig 1g-j). Next, we injected a retrograde monosynaptic virus into the Insula of AI9 dTomato mice (Fig 1h). We observed tomato-positive neurons in the contralateral Insula which are thus interhemispheric Insula neurons and represent homotopic labeling (Fig 1i-j). We quantified that 85.92 ± 2.64 % of cortical contralateral labeling was located in the homotopic cortical region and 14.08 ± 2.64 % in heterotopic cortical regions (Fig 1k-l). More precisely, we noted 44 % of homotopic labeling in the intermediate Insula, 43% in the posterior Insula, and 13 % in the anterior Insula (Supp Fig 1l). Insula interhemispheric neurons were mainly located in layer II/III (Supp Fig1m-n). We found 70.93 % of Insula interhemispheric neurons in layer II/III and 29.07 % in layer V/VI.

**Figure 1.**
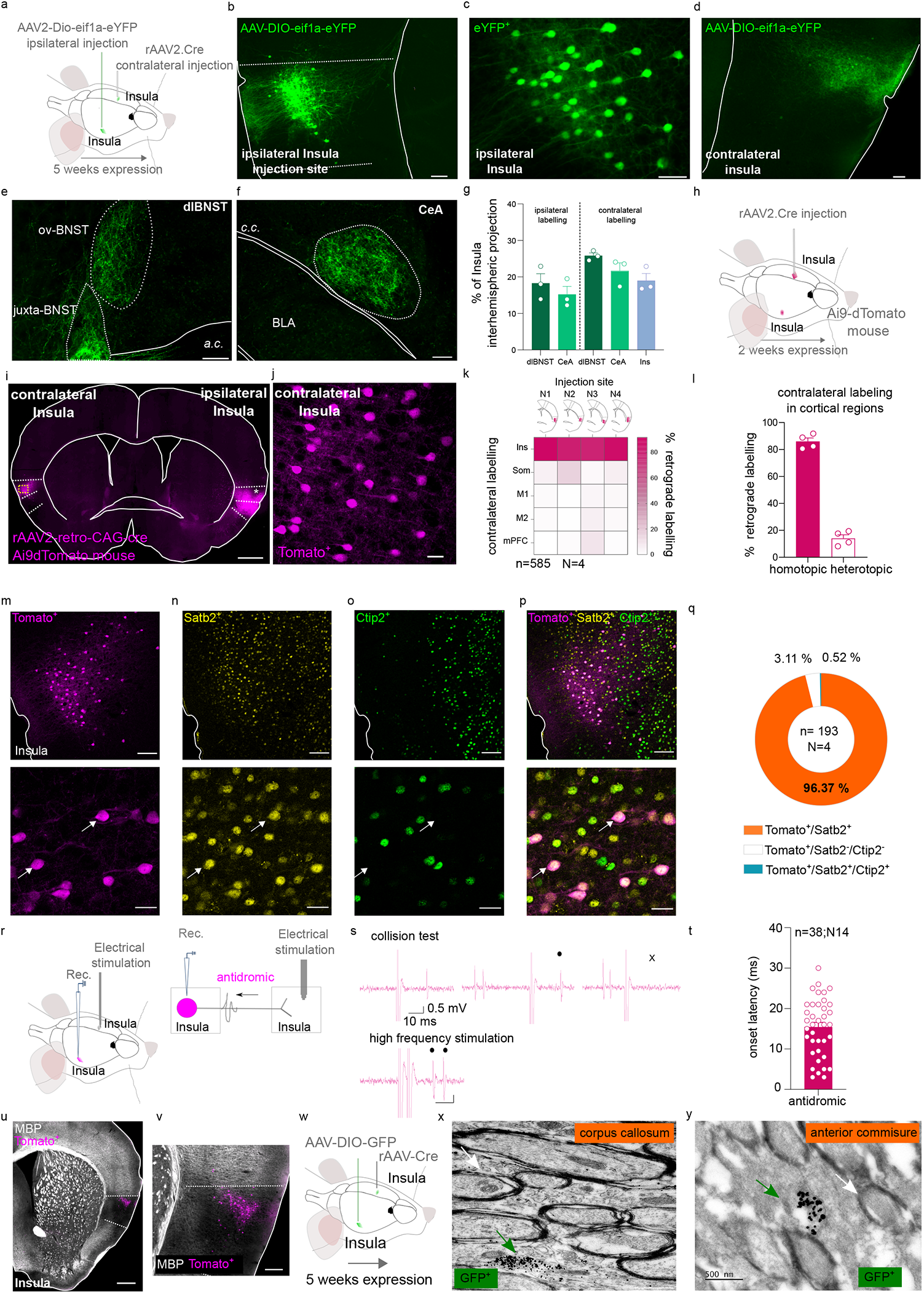
Anatomical, molecular, and electrophysiological characterization of Insula interhemispheric neurons. **a**.,**h**.,**r**.,**w**. Experimental design. **b-f**. Representative epifluorescent image of a coronal slice of brain injected with an AAV2-DIO-eif1a-eYFP anterograde virus in the Insula showing the injection site in the insula (b, c) and projections to the contralateral insula (d), the dlBNST (e), the CeA (f). Scale 100 µm for b,d,e,f and 50 µm for c. g. Quantification of the Insula interhemispheric projections in bilateral dlBNST, CeA, and contralateral Insula. **i**,**j**. Representative epifluorescent image of a coronal slice of brain injected with a rAAV2-retro-CAG-Cre retrograde virus in the ipsilateral insula in AI9dTomato mouse (i, scale 500 µm) with a high magnification of contralateral labeling in Insula (j, scale 25 µm). **k**,**l**. Quantification of contralateral labeling in Insula (homotopic labeling) and other cortical regions (heterotopic labeling). **m-p**. Immunofluorescence confocal images showing Insula interhemispheric neurons (tomato labeling, m), insula Satb2 staining (yellow labeling, n), insula Ctip2 labeling (green labeling, o), and the overlay at low (top, scale 100 µm) and high magnification (bottom, scale 25 µm). At the bottom, white arrows show examples of Tomato and Satb2 colocalizations. **q**. Quantification of Tomato, Satb2, and Ctip2 colocalization in the Insula. **s**. Representative traces showing a collision test and a high-frequency stimulation protocol for an Insula interhemispheric neuron projecting to the other Insula^55^. **t**. Histogram of the onset latency of Insula antidromic responses. **u**,**v**. Representative epifluorescent image at low (left, scale bar 500 µm) and high magnification (right, 150 µm) of MBP staining (grey labeling) and Insula interhemispheric neurons (tomato labeling). **x**,**y**. Representative image obtained with electron microscopy showing immunogold GFP labeling of unmyelinated interhemispheric Insula axons passing through the corpus callosum (x) or the anterior commissure (y) (green arrow). White arrow shows an example of a myelinated axon (scale 500 nm). *dlBNST: dorsolateral bed nucleus of the stria terminalis; ovBNST: oval-BNST; juxta-BNST: juxtacapsular BNST; a*.*c*.: *anterior commissure; cc*.*corpus callosum; BLA: basolateral amygdala; CeA: central amygdala; Ins: Insula; Som: Somatosensory cortex; M1: primary Motor cortex; M2:secondary Motor cortex; mPFC: medial Prefrontal cortex; Rec: recording; MBP: myelin basic protein*. n: number of neurons; N: number of mice.

We next molecularly characterized these Insula interhemispheric neurons that project to dlBNST and CeA. Interhemispheric neurons identified with tomato labeling specifically colocalized with Satb2 molecular marker without any colocalization with Ctip2, two transcriptional factors implicated in cortical development and maturation (Tomato^+^/Satb2^+^ colocalization: 96.13 ± 1.91%, Fig 1m-q). We next determined whether Insula interhemispheric neurons are exclusively pyramidal neurons or if they could be GABAergic projection neurons. No colocalization of Insula interhemispheric neurons with parvalbumin (PV) or glutamic acid decarboxylase (GAD67) staining was detected (Supp Fig 1o-q), thereby confirming that these Insula interhemispheric neurons belong to the category of excitatory pyramidal neurons.

By using *in vivo* electrophysiology in anesthetized mice, we functionally confirmed the reciprocal connectivity between both Insula (Fig 1r-s). Indeed, we recorded typical antidromic responses characterized by a collision test and, or high-frequency stimulation tests evoked by the electrical stimulation of their terminals in the contralateral Insula (Fig 1r, s). Interestingly, we observed a large variability in the latencies of antidromic responses (ranging from 3 to 30 ms; Fig 1t). One parameter influencing the action potential velocity conduction is the degree of myelination. Intriguingly, the Insula is a unique and specific cortical region that poorly expresses the myelin basic protein (MBP), an oligodendrocyte protein essential for myelin wrapping of axons in adult mice (Fig 1u-v). By using the double viral approach to identify Insula interhemispheric neurons (with GFP labeling) coupled with electron microscopy preparation, we showed that all Insula interhemispheric neurons observed had unmyelinated axons passing through the corpus callosum or the anterior commissure (Fig 1w-y;0 GFP^+^ myelinated axons out of n=59 GFP^+^ neurons, N=4 mice).

Thus, we highlight a novel neuronal subpopulation in the Insula that is the Insula interhemispheric pyramidal subpopulation characterized by bilateral projections to both dlBNST and CeA, mainly located in layer II/III, with a specific expression of the molecular marker Satb2^+^ and unmyelinated axons.

### Insula interhemispheric neurons provide a synaptic-excitatory drive on Insula interhemispheric circuit

We demonstrated that Insula interhemispheric neurons make asymmetric synapses, which are excitatory in function, with the contralateral Insula, the CeA, and the dlBNST (Fig2 a-d). Insula to CeA and to dlBNST synapses have been previously described ^25–29^. However, the Insula to Insula synapses remained poorly characterized. Studies that have functionally studied interhemispheric synaptic transmission have mainly studied the motor cortex and have demonstrated the importance of inhibition of the contralateral cortex in the execution of lateralized movements^30–32^. The Insula is involved in integrating of exteroceptive and interoceptive signals, contralateral inhibition does not seem necessary for the execution of emotional tasks. To complete our anatomical data, we tested the hypothesis that Insula stimulation may trigger an excitation in the contralateral Insula side by using *ex vivo* and *in vivo* electrophysiology in mice. First, we recorded contralateral Insula pyramidal neuron responses evoked by ipsilateral Insula optogenetic stimulation by using *ex vivo* electrophysiology (Fig 2e-g). We found that 87.5 % of the total recorded Insula pyramidal neurons respond by an excitation followed by inhibition to the ipsilateral Insula fiber optogenetic stimulation while 12.5% of these pyramidal neurons respond only by an excitation (Fig 2h-j; EPSC amplitude -267.9± 41.64 pA; IPSC amplitude 582.1 ± 125.9 pA). In addition, only inhibitory current is blocked by TTX+4AP pharmacological cocktail bath application suggesting that excitatory transmission is monosynaptic while inhibitory current is polysynaptic (Fig 2k; EPSC amplitude in aCSF: -242.8 ± 59.29 pA; EPSC amplitude in TTX+4AP: -247.3 ± 101.8 pA; IPSC amplitude in aCSF: 508.2 ± 123.4 pA; IPSC amplitude in TTX + 4AP: 5.167 ± 5.02 pA, IPSC amplitude aCSF vs TTX + 4AP: Wilcoxon test W=21,p=0.0313; EPSC amplitude aCSF vs TTX+4AP: Wilcoxon test W=1;p>0.05). IPSC response latency is also delayed compared to EPSC response latency (EPSC latency: 1.88 ± 0.22 ms; IPSC latency: 4.46 ± 0.25 ms, Mann-Whitney test U=2, p<0.0001). These data suggest that activation of Insula interhemispheric neurons drives monosynaptic excitation on Insula contralateral pyramidal neurons followed by polysynaptic feedforward inhibition.

**Figure 2.**
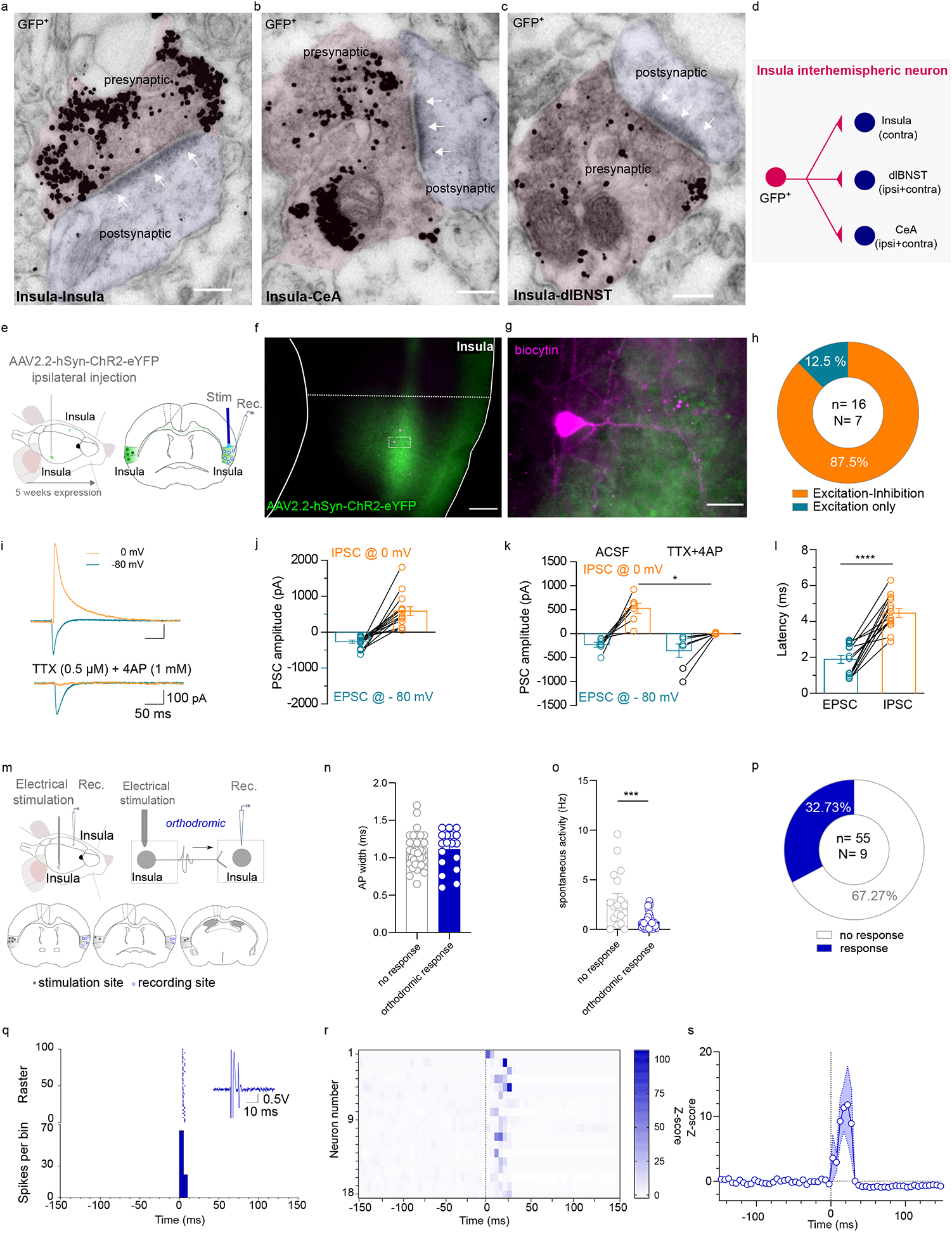
Functional characterization of insula interhemispheric circuit. **a-c**. Representative images of immunogold GFP labeling obtained with electron microscopy showing asymmetric synapses (at white arrows) for Insula to Insula synapses (a), Insula to CeA synapses (b), and Insula to dlBNST synapses (scale bar: 150 nm). **d**. Schematic representation of insula interhemispheric circuit. **e**. *Ex vivo* electrophysiological experimental design. **f**,**g**. Representative example of a histological control showing insula fibers expressing the Channelrhodhopsin (green labeling) projecting to the contralateral insula and insula recorded neurons filled with biocytin (tomato labeling) at low (f, scale bar: 150 µm) and high magnification (scale bar: 25 µm). **h**. Quantification of contralateral Insula pyramidal neuron responses to ipsilateral insula optogenetic stimulation. **i**. Representative traces of evoked ESPC recorded at -80 mV and IPSC recorded at 0mV in Insula pyramidal neuron before (top) and after TTX+4AP bath application (bottom). **j, k**. Group mean of evoked PSC amplitude of insula pyramidal neurons (j) and after TTX+4AP (k). **l**. Group mean of PSC response latency of insula pyramidal neurons. **m**. Experimental design (top) and cartography of stimulation and recording sites in the insula (bottom). **n**,**o**. Group mean of AP width (n) and spontaneous firing rate (o) of all the recorded insula neurons. **p**. Quantification of insula-responsive neurons to the electrical stimulation of the contralateral insula. **q**. Typical PSTH and raster show a contralateral Insula-evoked excitatory response of an Insula neuron. Electrical stimulus at 0 ms, with 5 ms bin width. **r**. Heatmap plot of Z-scored PSTH traces for each individual responsive Insula neuron to an Insula contralateral electrical stimulation. The electrical stimulus is represented by a vertical black line at 0 ms. **s**. Mean Z-score of PSTH over all responsive insula cells. Stim: stimulation; Rec: recording; PSC: postsynaptic current; EPSC: excitatory PSC; IPSC: inhibitory PSC; AP: action potential. n: number of neurons; N: number of mice.

Secondly, to decipher the net *in vivo* integrative effect of Insula interhemispheric transmission, we performed *in vivo* electrophysiology in anesthetized mice (Fig 2m). We observed that 32.73 % of all contralateral Insula recorded neurons respond to ipsilateral Insula electrical stimulation (Fig 2p). These insula-responsive neurons are characterized by a half-action potential width of 1.08 ± 0.03 ms and a spontaneous firing frequency of 0.78 ± 0.13 Hz (Fig 2n-o). Insula interhemispheric neuronal stimulation triggered excitatory responses on the contralateral Insula neurons with 10.61 ± 1.715 ms response latency (Fig 2q-s).

Together, these results suggest that interhemispheric neurons contact both excitatory and inhibitory contralateral insula neurons, and feed-forward inhibition was activated within ∼2.5 ms after the onset of excitation in both cell types, creating a precise temporal excitation in the Insula network.

### Genetic selective ablation of Insula interhemispheric communication disrupts social preference following acute social isolation

Lastly, to determine whether Insula interhemispheric communication plays a role in social interaction and anxiety processing, we measured mouse social interaction with a three-chamber social test in two different housing conditions associated with a caspase viral approach strategy to selectively lesion Insula interhemispheric neurons, leading to split Insula mice (Fig 3a-b). Since rodents are innately pro-social species, social isolation represents an aversive experience. Previous studies have shown that structures involved in the Insula interhemispheric network are recruited and display plastic adaptive neuronal responses after acute isolation such as the dorsal raphe nucleus or the dlBNST^33,34^. To elucidate whether Insula interhemispheric neurons are crucial to developing adaptive social behavior after this aversive event, we assessed social preference in group-housed conditions and 24 h after acute social isolation in the control group mice and the caspase group. We injected a retrograde monosynaptic virus (rAAV2-retro-CAG-Cre) in two main outputs of the insula interhemispheric neurons (contralateral Insula and ipsilateral CeA) and an AAV-Flex-taCaspase-TEVp or the control virus (AAV-Flex-eGFP) in the ipsilateral Insula (Fig 3a). We first confirmed that the caspase viral strategy specifically lesioned Insula interhemispheric neurons as illustrated in the histological control example and quantified by NeuN fluorescence density in layer II/III of Insula (Fig 3c-e). NeuN fluorescence density is specifically decreased in the layer II/III of the Insula caspase injection site compared to its contralateral Insula control site (Insula control site NeuN density: 44.15 ± 2.37; Insula caspase injection site NeuN density: 34.01± 2.86, Two-tailed Paired-t-Test, t(8)=2.788, p=0.0236) without altering other proximal cortical regions as the somatosensory cortex (NeuN density in Somatosensory cortex control site: 31.28 ± 2.33; NeuN density in the other Somatosensory cortex site: 33.96 ± 0.91, Two-tailed paired t-Test t(8)=1.101, p>0.05). Under the group-housed condition, both control and caspase mice spent more time around the social enclosure compared to the object enclosure without differences in the three-chamber test (Fig 3f, time spent around for ctrl: object 65.2 ± 7.4 s vs social 149.4 ± 9.6 s; caspase: object 57.14 ± 5.9 s vs social 144.9 ± 10.72 s; Two Way repeated measure Anova, zone x virus interaction effect F(1.18)=0.02973, p>0.05; zone main effect F(1.18)=67.17,p<0.0001; virus main effect, F (1.18)=0.9348, p>0.05 ). Control and caspase mice developed a social preference in the group-housed condition indicated by a social preference ratio higher than 0.5 (Fig 3g; ctrl social preference ratio: 0.69 ± 0.04 and caspase social preference ratio 0.72 ± 0.02; ctrl vs caspase social preference ratio, Mann-Whitney U=47, p>0.05; One sample Wilcoxon test for ctrl: W=66, p=0.001, and caspase: W=45, p=0.0039). They spent a similar amount of time in the social zone (Fig 3h-i; time in the social zone for ctrl: 49.79 ± 3.2 %; for caspase: 48.31 ± 3.57 %; two-tailed unpaired t-test, t(18)=0.3089, p>0.05; mean social bout duration for ctrl: 9.4 ± 0.71%; for caspase: 8.13 ± 0.78 %; two-tailed unpaired t-test, t(18)=1.197).

**Figure 3.**
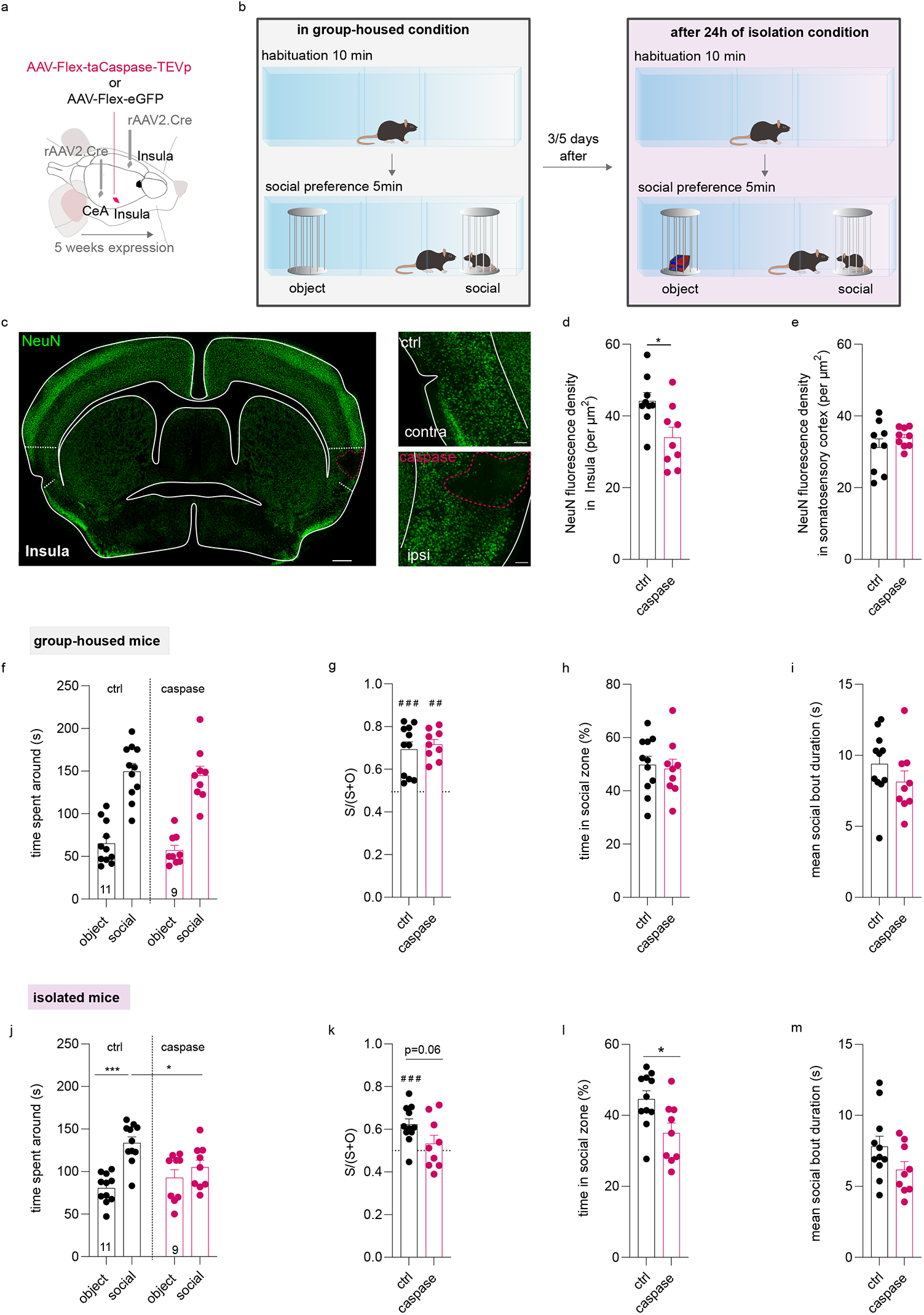
Split insula interhemispheric communication disrupts social preference only after acute social isolation. **a**,**b**. Experimental design for the viral injection (a) and behavioral assay (b). **c**. Example of a histological control of caspase lesion in the insula identified by NeuN immunofluorescence labeling (green labeling) taken at epifluorescence microscope at low (left, scale bar: 500 µm) and high magnification (right, scale bar: 100 µm). **d.e**. Quantification of NeuN fluorescence density in the control side (contralateral side to the injected lesion side) compared to the caspase injection side in the Insula cortex (d) and in the Somatosensory cortex (e). **f**,**j**. Quantification of the time spent around the object and social enclosures in the three-chamber test in grouped housed condition (f) or after acute social isolation (j) in insula control and insula caspase groups. **g**,**k**. Social preference ratio in the control group and caspase group in group-housed mice (g) or isolated mice (k). **h**,**l**. Time spent in the social zone between control and caspase mice in group-housed (h) or acute isolated condition (l). **i**,**m**. Mean social bout duration in control and caspase group in group-housed (i) and acute isolated housing condition (m). *ctrl: control, S: time in the social zone; O: time in the object zone; contra: contralateral; ipsi: ipsilateral*.

We next tested how social isolation affects attention to social stimuli in split Insula mice. After acute social isolation, ctrl mice spent more time around the social enclosure compared to the object enclosure while caspase mice spent the same time around both enclosures (Fig 3j, time spent around for ctrl: object 80.49 ± 5.25 s vs social 133.7 ± 7 s; caspase: object 92.91 ± 9.31s vs social 105± 8.51 s; Two-Way repeated measure Anova, Zone x virus interaction effect, F(1.18)=5.011, p=0.0381; zone main effect F(1.18)=12.65, p=0.0023; virus main effect F(1.18)=0.3213, p>0.05; Bonferroni *post hoc* test ctrl social caspase social p=0.02; ctrl social vs control object: p=0.0008 ). Control group mice still presented social preference after acute isolation whereas caspase mice did not, despite significant differences between groups (Fig 3k; ctrl social preference ratio: 0.62 ± 0.03 and caspase social preference ratio 0.53 ± 0.04; Two-tailed unpaired t-test, t(18)=1.956, p=0.0662; One sample t-test for ctrl: t(10)=4.69, p=0.0009; for caspase: t(8)=0.8273, p>0.05). Caspase mice spent less time in the social zone compared to control mice only after acute social isolation (Fig 3l; time in the social zone for ctrl: 44.58 ± 2.33 %; for caspase: 35.01 ± 2.84 %; Two-tailed Unpaired t-Test, t(18)=2.632, p=0.0169). There was no difference in the mean social bout duration between groups (Fig 3m; mean social bout duration for ctrl: 7.8 ± 0.72 s; for caspase: 6.17 ± 0.58 s; Two-tailed unpaired t-test, t(18)=1.707, p>0.05).

Insula is activated during anxious situations and Insula overactivation has been detected in patients with Anxiety disorders ^11,14,35–39^. Since Insula interhemispheric neurons project to CeA and dlBNST, we next investigated the impact of Insula interhemispheric communication split on unconditioned anxiety tests in mice. In rodents, anxiety can be measured based on the innate approach/avoidance behaviour in a novel environment. We didn’t detect a change in the time spent and the number of visits in the center of the open field, nor in the total distance travelled (Supp Fig 2 a-d, Time spent in the center for ctrl: 16.65 ± 2.06 %; caspase: 17.47 ± 1.45 %, two-tailed unpaired t-test, t(18)=0.3088, p>0.05; the number of visits in the center for ctrl: 57.45 ± 5.42 visits; caspase: 62.89 ± 4.26 visits, two-tailed unpaired t-test, t(18)=0.7612, p>0.05; total distance travelled for ctrl: 4367 ± 309.3 cm; caspase: 4526 ± 214.2 cm, two-tailed unpaired t-test, t(18)=0.404, p>0.05 ). In addition, we didn’t observe a difference in the time spent and the number of entries in the open arms in the elevated plus maze nor in the total distance travelled between control and caspase mice (Supp Fig 2 e-h, Time spent in the open arms, ctrl: 4.47 ± 1.01 %; caspase: 5.24 ± 1.37 %, Mann-Whitney, u=41.5,p>0.05; the number of entries in the OA for ctrl: 4.091 ± 0.72 visits; caspase: 3.89 ± 0.89 visits, Two-tailed Unpaired-t-test, t(18)=0.1788, p>0.05; total distance travelled for ctrl: 967 ± 80.99 cm; caspase: 1117 ± 69.14 cm, Two-tailed unpaired t-test, t(18)=1.375,p>0.05 ).

These data show that Insula interhemispheric communication split leads to impairment of social preference only after acute social isolation without interfering with anxiety-like behaviors.

## Discussion

We unraveled the anatomical and molecular phenotype of an interhemispheric neuronal subpopulation in the Insula that belongs to a restricted network enrolling both bilateral dlBNST/CeA and contralateral Insula, mainly located in the layer II/III, characterized by unmyelinated axons and expressing the transcriptional factor Satb2^+^. Our findings enlightened the contribution of the Insula interhemispheric neurons in social processing. Selective ablation of Insula interhemispheric neurons leads to a reduced interest in social stimulus after acute social isolation which is a maladaptive behavior. This data suggests that the Insula interhemispheric communication split created an imbalance in social homeostasis processes.

Pioneering studies including lesioning approaches and split-brain patient cases who presented surgical callosal incisions shed light on brain lateralized functions^40,41^. One of the most studied cases of cortical interhemispheric communication has been described in the motor cortex region. For instance, the execution of lateralized motor movement requires inhibition of the contralateral side induced by interhemispheric cortical inhibition^30^. The degree of myelinization of axons directly impacts the efficiency of interhemispheric communication. Decreased efficiency in interhemispheric inhibition has been observed in children who are characterized by a hypo-myelination of callosal neurons resulting in difficulties in generating unilateral motor movement and leading to non-lateralized mirror movements^42^. The nature and the recruitment of cortical interhemispheric communication may depend on the type and the complexity of the performed task as well as the cortex involved^43^. Despite mammalian evolution, communication between two brain hemispheres presents some similarities across species between rodents, non-human primates, and humans. Here, we found that stimulation of Insula interhemispheric neurons leads to excitation of the Insula contralateral side in mice (Fig 2h-s).

In general, myelination of axons which is a dynamic process ensures a fast and precise transfer of information to the targeted zone. Thus, the degree of myelination is one of the parameters that influence the conduction velocity and define the efficiency of neuronal communication. Unexpectedly, the entire population of interhemispheric neurons in the insular cortex are unmyelinated neurons in adult mice under physiological conditions (Fig 1x, y). We confirmed this phenomenon by a low density in MBP immunostaining in the Insula (Fig 1u,v) contrary to what has been described in other cortical regions^44 45^. This lack of myelination on Insula interhemispheric axons may explain the variability in their onset latency in response to axonal stimulation (Fig 1t). Future studies would be required to understand this atypical Insula signature and the potential implication of myelination process within Insula interhemispheric neurons at synaptic, circuit and behavioural levels in physiological and pathological states.

In this study, we found that selective ablation of Insula interhemispheric neurons impaired social preference only following acute social isolation (Fig3 j-l). After this aversive event, control mice developed adaptive behavior that favours social interactions compared to object interactions which restore social homeostasis. Caspase mice did not present this appropriate strategy after acute social isolation which can suggest reduced attention toward social stimulus and/or a lack of motivation for orienting to social cues. Interestingly, a clinical study monitoring the Insula interhemispheric communication evoked by the presentation of social versus non-social stimuli in children with neurotypical development or with ASD, reported that only children with ASD presented an hyperconnectivity of this pathway^12^. Together, these results suggest that Insula interhemispheric communication may be recruited when social homeostasis is unbalanced to promote adaptive and appropriately motivated behavior to seek for social contacts. Here, we demonstrated that the interhemispheric neurons occupy a privileged position in synchronizing the activity of the Insula/CeA/dlBNST network in the two hemispheres. Interestingly, interhemispheric neurons of the Insula and dopaminergic neurons of the Dorsal Raphe Nucleus also project massively to the dlBNST and the CeA^46^. Future studies need to elucidate how the dopamine/glutamate interplay controls the CeA/dlBNST network. Thus, all of these data suggest that interhemispheric neurons in the Insula play a specific role for social processing. This specificity is confirmed because we did not detect an anxiety phenotype after Insula interhemispheric deletion by using an open field or an elevated plus maze (Supp Fig2).

Here, we assessed a right unilateral lesion of Insula interhemispheric neurons by using a double viral approach (Fig 3a). One limitation of our study is that our viral strategy may minimize the effect observed by manipulating a smaller quantity of cells. However, to be selective of Insula interhemispheric neurons without targeting interneurons (Supp Fig 1o-q), we restricted our ablation to the right Insula interhemispheric neurons. Despite this limitation, our viral approach allowed a selective ablation of right interhemispheric neurons in the Insula which can be useful for future investigations of Insula lateralization. Indeed, a lateralization of Insula with a right dominance has been observed with cFos analysis in response to intraperitoneal injection of lithium chloride, an aversive visceral stimulus or in feeding behavior^47,48^. Additional studies would be necessary to define the Insula laterization processes. This study has been carried on adult male mice and future detailed analysis will be valuable to dissect age- and sex-dependent effects.

Our study, by demonstrating the role played by the interhemispheric neurons of the Insula in social processing, reinforces the concept of cellular diversity of the Insula^27,37,45,48–52^. Another avenue for future research motivated by the present study will be to examine the development, maturation and neuromodulation of this interhemispheric Insula circuit in physiological and pathological states where social processing is altered as in the ASD mouse model.

## Author contributions

C.G. and F.G conceived and designed the experiments. C.G. performed and analyzed the anatomical study with the participation of E.L. and M.G. E.D., E.B. performed the data with electron microscopy. C.G and A.G. performed and analyzed the *in vivo* electrophysiology in anesthetized mice. A.T and J.B. designed, performed, and analyzed the *ex vivo* electrophysiological experiments. C.G. performed and analyzed the behavioural study. C.G. and F.G. wrote the manuscript and C.G. prepared the figures.

## Acknowledgments

C.G. is supported by the Foundation for Medical Research: ARF20170938746. This work was supported by recurrent funding from the University of Bordeaux and the CNRS. This study also received financial support from the French government in the framework of the University of Bordeaux’s IdEx “Investments for the Future” program / GPR BRAIN_2030. The microscopy was done in the Bordeaux Imaging Center a service unit of the CNRS-INSERM and Bordeaux University, member of the national infrastructure France BioImaging supported by the French National Research Agency (ANR-10-INBS-04). The help of Sebastien Marais is acknowledged.

## Conflict of interests

The authors declare no conflict of interest.

## STAR Methods

### Key resource table

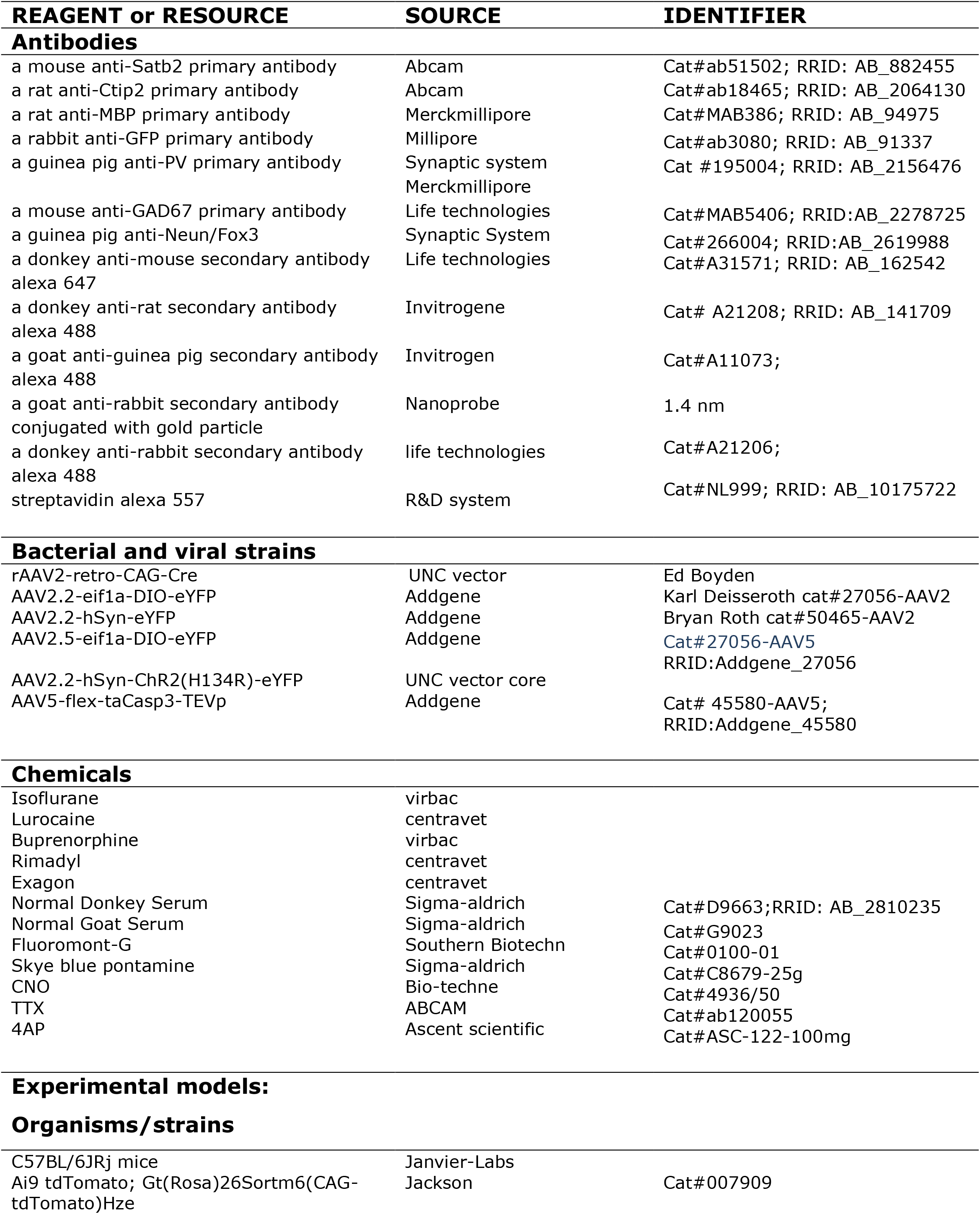
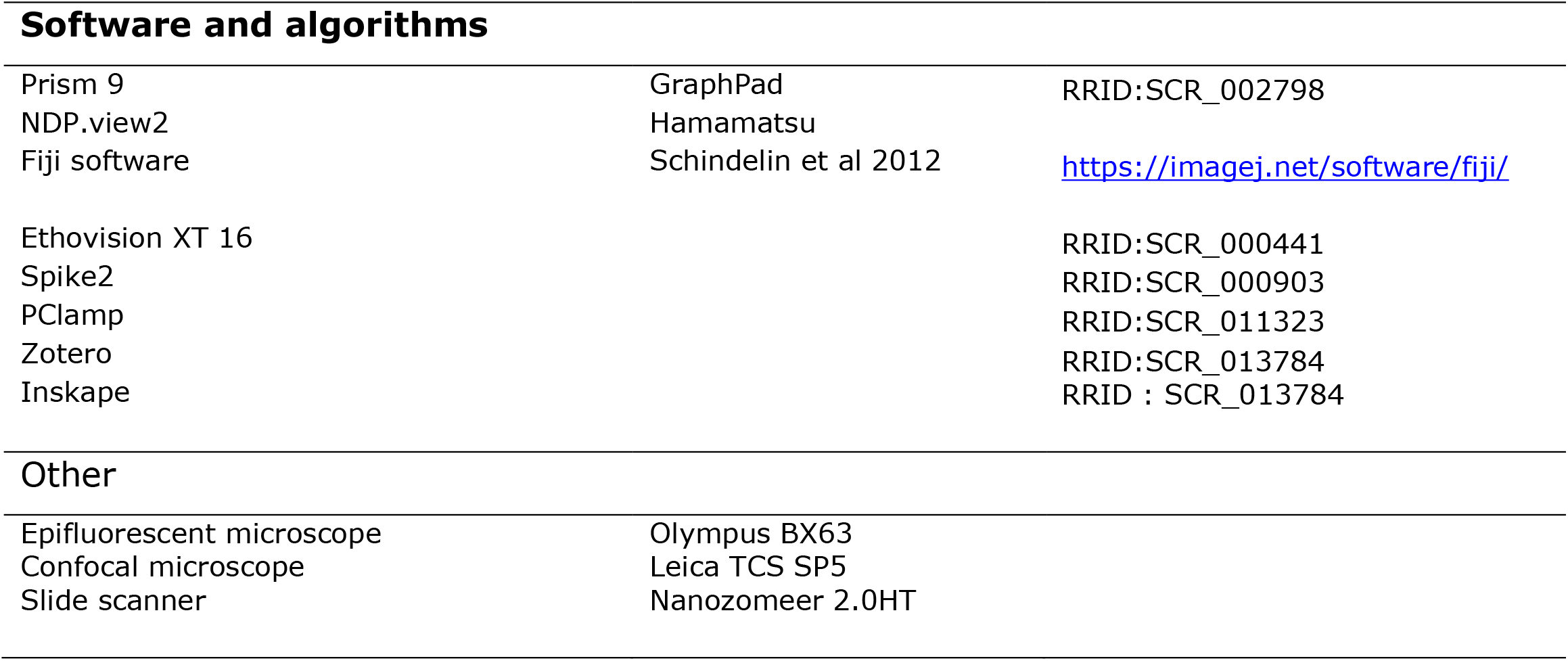

### Resource availability

#### Lead contact

Further information and requests for resources and reagents should be directed and will be fulfilled by the lead contact, François Georges.

### Data and code availability

This paper does not report original code.

### Materials availability

This study did not generate new unique reagents.

## Experimental model and subject details

### Animals

Male C57BL/6JRj (≥ 10 week old; Elevage Janvier, France) were used. Male Ai9 tdTomato also called as Gt(Rosa)26Sortm6(CAG-tdTomato)Hze (stock number 007909, from Jackson; C57BL6/j genetic background) were also used. Mice were housed three to five per cage under controlled conditions (22-23°C, 40 % relative humidity, 12 h light/dark illumination cycle; with lights on at 07:00). Mice were acclimatized to laboratory conditions at least one week prior to experiments, with food and water *ad libidum*. All procedures were conducted in accordance with European directive 2010-63-EU and with approval from the Bordeaux University Animal Care and Use Committee (license authorization 21134).

### Methods details

#### Viruses and Drugs

rAAV2-retro-CAG-Cre (2,8x10^12^ vg/mL; UNC Vector Core, Boyden);AAV2.2-eif1a-DIO-eYFP (3x10^12^ vg/mL; Addgene); AAV2.5-eif1a-DIO-eYFP (1x10^13^ vg/mL; Addgene);AAV2.2-hSyn-eYFP (3x10^12^ vg/mL; 50465-AAV2,Addgene); AAV2.5—eif1a-DIO-eYFP (1x10^13^ vg/mL; 27056-AAV5, Addgene); AAV2.2-hSyn-ChR2(H134R)-eYFP (3.1x10^12^ vg/mL; UNC, AV4384G); AAV5-flex-taCasp3-TEVp (7x10^12^ vg/mL; Addgene); Tetrodotoxin (TTX, 0.5 µM, abcam ab120055); 4 aminopyridine (4AP; 1 mM, ascent scientific, asc-122-100mg)

#### Surgery

Stereotaxic surgery for anatomy, *ex vivo* and *in vivo* electrophysiology experiments, and behavioral tests were performed under a mixture of isoflurane and oxygen as previously described^53^. Mice were placed on a stereotaxic frame and received a subcutaneous dose of buprenorphine (0.1 mg/kg, except for *in vivo* electrophysiology experiments) and local injection of an analgesic prior to skin incision (lurocaine, 7mg/kg). Single or bilateral craniotomy was made over the insular cortex at the following coordinates (+0.14 mm/bregma, ±3.8 mm/midline, 2.2 mm/brain surface), the CeA (-1.58 mm/bregma, +2.4 mm/midline, 3.9 mm/brain surface). Viruses were injected via a glass micropipette into the region of interest. Following injections, the incision was closed with sutures and mice were let to wake up on a heating plate. For all the experiments the virus was incubated at least four weeks before proceeding with further manipulation except for the experiment with the retrograde virus (rAAV2-retro-CAG-cre) injection in AI9 dtTomato in which only two weeks were sufficient to clearly identify reporter protein expression.

#### Immunohistochemistry

Mice were deeply anesthetized with a mixture of isoflurane and oxygen and received an i.p. lethal dose of exagon (300 mg/kg) and lidocaine (30mg/kg). Mice were perfused transcardially with phosphate-buffered saline (PBS 1X) and incubated (48h/4°C) in 4% paraformaldehyde. Coronal slices were cut at 50 μm and washed three times in PBS 1X before incubation in the blocking solution containing 0.03% Triton X-100 and 10% donkey serum or goat serum. Sections were incubated (overnight per 4°C) with a mouse anti-Satb2 primary antibody (1/300; abcam ab51502), a rat anti-Ctip2 primary antibody (1/500; Abcam ab18465), or with a rat anti-MBP (1/500,Merckmillipore), a guinea pig anti-NeuN/Fox3 (1/1000,cat 26604, Synaptic system), a rabbit anti-GFP primary antibody (1/1000; Millipore, AB3080), a mouse anti-GAD67 primary antibody (1/500; Millipore MAB5406), an guinea pig anti-PV primary antibody (1/1000, synaptic system, cat#195004). After washing sections were incubated overnight at 4° C with a donkey anti-mouse secondary antibody (labeling of Satb2, 1/500, life technologies A31571, alexa 647), a donkey anti-rat secondary antibody (labeling of Ctip2 or labeling of MBP, 1/500, life technologies A21209, alexa 488), a goat anti-mouse secondary antibody (labeling of GAD67,1/500, Invitrogene A21202, alexa 488), a goat anti-guinea pig secondary antibody (labeling of PV or Neun/Fox3,1/500, Invitrogen A11073, alexa 488), a donkey anti-rabbit (labelling GFP,1/500,life technologies A21206, alexa 488), streptavidine (labeling of biocytin, R&D system NL 999, 1/500, alexa 557). Sections were washed and then mounted in Fluoromont-G medium (Southern Biotech), coverslipped, and imaged on a fluorescent microscope as a confocal microscope (Leica SP5) or a slide scanner (Nanozomeer 2.0HT), or an epifluorescent microscope (Olympus BX63). Photomicrographs were taken and displayed using image J to adjust the contrast and or perform Z stack images.

### Electron microscopy sample preparation

#### Tissue preparation

Mice were deeply anesthetized and perfused transcardially with a mixture of 3% paraformaldehyde (PFA) and 0.5% glutaraldehyde in 0.1M phosphate buffer at Ph 7.4. Brains were quickly removed, left overnight in 3% PFA at 4°C. Coronal sections of the brain were cut on a vibrating microtome at 50 µm, collected in PBS, cryoprotected, freeze-thawed, and stored in PBS with 0.03% sodium azide until use.

#### Immunogold experiments

GFP was analysed at electron microscopic level in Insula, Corpus Callosum, Anterior Commissure,dlBNST and CeA. GFP was detected by the preembedding immunogold technique, sections were incubated in 4% NGS for 45 min and then in a mixture of a rabbit anti-GFP (1/5000) antibody supplemented with 1% NGS overnight at RT. After washing, in PBS and PBS-BSAc (aurion, the Netherlands), the sections were incubated for 3 hours at RT in Goat anti-rabbit IgG conjugated to ultrasmall gold particles (1.4nm; nanoprobes) diluted 1/100 in PBS-BSAc-gel. The sections were washed and post-fixed in 1% glutaraldehyde in PBS for 10 min. After washing in PBS and water distilled, the immunogold signal was intensified using a silver enhancement kit (HQ silver; Nanoprobes, Yaphank, NY) for 8 min at RT in the dark. After several washes in PBS, the sections were then processed for electron microscopy.

The sections were post-fixed in 0.5% osmium tetroxide and dehydrated in ascending series of ethanol dilutions that also included 70% ethanol containing 1% uranyl acetate. The sections were post-fixed, dehydrated, and included in resin ( Durcupan ACM; Fluka). Serial ultrathin sections were cut with a Reichert Ultracut S, contrasted with lead citrate and imaged in a transmission electron microscope (H7650, Hitachi) equipped with a 467 SC1000 Orius camera (Gatan).

### *Ex vivo* Electrophysiology

After allowing at least 4 weeks for viral vector expression acute coronal brain slices containing the Insula were cut on a vibratome (VT1200S; Leica microsystems). Mice were deeply anaesthetized by i.p. injection of a mixture of ketamine-xylazine (100mg/kg and 20mg/Kg, respectively). A thoracotomy followed by a transcardiac perfusion with a saturated (95%O_2_ / 5%CO_2_), iced-cold solution (cutting solution) containing 250 mM sucrose, 10 mM MgSO_4_·7H_2_O, 2.5 mM KCl, 1.25 mM NaH_2_PO_4_·H_2_O, 0.5 mM CaCl_2_·H_2_O, 1.3 mM MgCl_2_, 26 mM NaHCO_3_, and 10 mM D-glucose (pH 7.4) was performed. The brain was then quickly removed from the skull, blocked in the coronal plan, glued on the stage of the vibratome, submerged in iced-cold, saturated cutting solution and cut in 300-µm thick sections. Brain slices were transferred in a storage chamber at 34°C for 1 h in an artificial cerebral spinal solution (referred as « recording ACSF ») saturated by bubbling 95%O_2_ / 5%CO_2_ and containing 126 mM NaCl, 2.5 mM KCl, 1.25 mM NaH_2_PO_4_·H_2_O, 2 mM CaCl_2_·H_2_O, 2 mM MgSO_4_·7H_2_O, 26 mM NaHCO_3_, and 10 mM D-glucose, supplemented with 5 mM glutathion and 1 mM sodium pyruvate (pH: 7.4; Osmolarity : 310-315 mOsm). They were then maintained at room temperature in the same solution until recording.

Whole-cell patch-clamp recordings were performed in a submerged chamber under an upright microscope (AxioExaminer Z1; Zeiss) equipped with IR-DIC illumination. Slices were bathed in recording solution. Recording pipettes (5-7 MΩ) were prepared from borosilicate glass capillaries (GC150F-10; Harvard Apparatus) with a horizontal puller (Sutter Instrument, Model P-97). They were filled an internal solution composed of 135 mM K-gluconate, 3.8 mM NaCl, 1 mM MgCl_2_·6H_2_O, 10 mM HEPES, 0.1 mM Na_4_EGTA, 0.4 mM Na_2_GTP, 2 mM Mg_1.5_ATP, 5 mM QX-314 and 5 mM Biocytin (pH :7.25; Osmolarity: 290-295 mOsm). Experiments were conducted using a Multiclamp 700B amplifier and Digidata 1440 digitizer controlled by Clampex 10.6 (Molecular Devices) at 34°C. Data were acquired at 20 kHz and low-pass filtered at 4 kHz. Insula pyramidal neurons were visualized under IR-DIC microscopy and recognized by the triangular shape of their soma. All the recordings were performed in voltage-clamp mode at -80 and 0 mV to record light-evoked glutamatergic EPSC and GABAergic IPSC, respectively. Voltages were corrected off line for liquid junction potentials. Optical stimulations were achieved using a 473 nm diode pumped solid state laser (Optotronics, USA) connected to a 800 μm diameter optical fiber (Errol, Paris, France) positioned just above the surface of the slice next to the recording site. At the end of the day, brain slices were fixed in 4% PFA overnight and stored in 0.2% sodium azide-PBS until histological processing.

### *In vivo* electrophysiology

Electrical stimulation of the Insula. Bipolar electrical stimulation of the Insula was conducted with a concentric electrode (Phymep) and a stimulator isolator (800 µs, 0.2-1.8 mA; Digitimer).

#### Insula recordings

A glass micropipette filled with 2% pontamine sky blue solution in 0.5 M sodium acetate was lowered in the insula. The *in vivo* single-unit recordings were performed as previously described (Glangetas et al 2015). Briefly, the extracellular potential was recorded with an Axoclamp-2B amplifier and filter (300 Hz/0.5 kHz). Single neuron spikes were collected online (CED 1401, SPIKE 2; Cambridge Electronic Design). During electrical stimulation of one insula, cumulative peristimulus time histograms (PSTH) (5 ms bin width) of the contralateral Insula were generated for each neuron recorded. Electrical stimulation of the contralateral Insula was also used to test for antidromic activation of ipsilateral insula neurons using high-frequency stimulation and collision methods as previously described^54^. Driven impulses were considered antidromic if they met the following criteria: (1) constant latency of spike response (fixed jitter), (2) driven by each of the paired stimulus pulses at frequencies of 100 Hz or greater, and (3) collision of driven spikes by spontaneous impulses.

#### Histological control

At the end of each recording experiment, the recording pipette placement was marked with an iontophoretic deposit of pontamine sky blue dye (−20 μA; 30 min). To mark the electrical stimulation sites, +50 μA was passed through the stimulation electrode for 90 s. Then, mice were perfused with PBS 1x and stored for 48 h in PFA 4% at 4°C.

### Behavioral procedures

One week prior behavioral experiment, mice were progressively handled by the experimenter. For each behavioral test, mice were acclimatized at least 30 min in the experimental room. Between each mouse, the behavioral apparatus was cleaned with 70% ethanol and then water and dried between each test.

### Open Field test

Mice were placed in the corner of a square open field (40 × 40 cm) and were allowed to freely explore the open field for a 10-min period in 70 lux illumination conditions. Total distance travelled, velocity, and time spent in the zone during the session were automatically reported (Ethovision, Noldus).

### Elevated plus Maze

The elevated plus maze consisted of a platform of four opposite arms (30 cm x5cm) two of them are open and two are closed arms (enclosed by 25 cm high walls). The apparatus was elevated from the floor. The task was analyzed with the software Ethovision (Noldus) and we measured the time spent in each arm in trials of 10 min. The luminosity of the open arms was around 120 lux.

### Social preference test

A three-chamber rectangular plexigas arena (60x42x22 cm, Imetronic) divided into three chambers of the same dimension was used for this test. Briefly, each mouse was placed in the center of the arena and allow to freely explore the entire arena for a 10 min habituation period under approximatively 90 lux illumination condition. At the end of the habituation, the mouse was placed in the center of the arena, and two metallic enclosures (9 cm x 9 cm x 10 cm) were positioned in the center of the two outer chambers. One enclosure contained a juvenile unfamiliar mouse whereas the other enclosure was empty (inanimate object) for group-housed condition or filled with lego toys for isolated condition. The position of the two enclosures was counterbalanced to avoid any bias. The juvenile mice were previously habituated to the enclosure and the arena for a brief period of 2 days preceding the experiment with a 10-min session per day. The experimental mouse was allowed to freely explore the three-chamber arena for a 5 min period session. The time spent around the enclosures were manually scored. The stimulus interaction was scored when the nose of the experimental mouse was in closed proximity to the enclosure (approximatively around 2 cm).

### Data analysis

For *in vivo* electrophysiological experiments, cumulative PSTHs of insula activity were generated during stimulation of the contralateral insula. Excitatory magnitudes were normalized for different levels of baseline impulse activity. Baseline activity was calculated on each PSTH, during the 500 ms preceding the stimulation to generate a Z-score for each responding neuron.

#### For immunolabeling quantification

To quantify retrograde labelling (rAAV2-retro-CAG-cre/Ai9dtomato mouse), we acquired 3 slices for each Insula level (antero, intermediate and posterior level) per mouse, on a total of 4 mice with confocal microscope. For the co-localization of Tomato^+^ neurons with Satb2^+^ and Ctip2^+^ labelling, we took 3 pictures with confocal microscope per slice, on 3 slices per mouse, with a total of 4 mice and analysed co-localization on focal plan. For caspase lesion, we took one picture in the mid-Insula level with a slide scanner per mouse and quantify with Image J, the fluorescence density between contralateral (ctrl side) and ipsilateral lesion side (caspase side).

### Statistical analysis

Statistical outliers were identified with the Rout Q method (Q=1%) and excluded from the analysis. Normality was checked with the Shapiro-Wilk criterion and when violated, non-parametric statistics were applied (Mann–Whitney and Kruskal– Wallis, Wilcoxon test). When samples were normally distributed, data were analyzed with independent or paired one or two-tailed samples t-tests, one-way, two-way, or repeated measures ANOVA followed if significant by Bonferroni *post hoc* tests. Data are represented as the mean ± SEM, and the significance was set at *p* < 0.05. Data were analyzed using GraphPad Prism 9.

## Supplementary figure legends

**Supplementary Figure1. a**. Experimental design. Representative epifluorescent image of a coronal slice of brain injected with an AAV2-hSyn-eYFP anterograde virus in the insula showing the injection site in the Insula (b) and projections to the dlBNST (c), the CeA (d), the contralateral insula (e). Scale bar: 1 mm for (b) and 500 µm for (c-e). **f**. Quantification of the insula bilateral projections. g. Experimental design. **h-j**. Representative epifluorescent image of a coronal slice of brain injected with a rAAV2-Cre retrograde virus in the CeA coupled with an AAV5-DIO-eif1a-eYFP anterograde virus in the Insula showing the injection site in the insula (h) and projections to the dlBNST (h), the CeA (i), the contralateral insula (h,j).Scale bar: 500 µm (h, i) and 25 µm (j). **k**. Cartography of insula subregions. **l-n**. Quantification of contralateral labeling in the homotopic region in insula subregions (l), and layers (m,n) after rAAV2-retro-CAG-Cre retrograde virus injection in the insula cortex in Ai9 dTomato. **o-q**. Representative confocal images showing an absence of co-localization between PV (o, green) or GAD67 (p, green) immunofluorescence staining and interhemispheric insula neurons (tomato labeling and its quantification (q). scale bar: 25 µm. *ovBNST: oval-BNST; juxta-BNST: juxtacapsular BNST, am-BNST: anteromedial BNST; dlBNST: dorsolateral BNST, CeA: central amygdala, BLA: Basolateral amygdala; a*.*c*.: *anterior commissure; cc: corpus callosum; Claust: claustrum; GI/DI: granular/disgranular insula; AI: agranular insula; Nac: nucleus accumbens, mPFC: medial Prefrontal cortex; M1: primary Motor cortex; M2: secondary Motor cortex; PAG: periaqueductal grey substance; d: dorsal; l: lateral; v: ventral; PV: parvalbumin; GAD67: glutamic acid decarboxylase*.

**Supplementary Figure2. Caspase insula lesion does not modify locomotion or anxiety phenotype. a**., **e**. Example heatmaps of insula control and caspase mice performance in the open field test (a) or in the elevated plus maze (e). **b-d**. Quantification of the time spent in the center of the open field (b), the number of visits to the center (c), and the total distance traveled (d) in control and caspase mice. **f-h**. Quantification of the time spent and the number of entries in the open arms of the elevated plus maze, (f and g respectively) and the total distance traveled (h) in both groups. *Ctrl: control; OA: open arms; CA: closed arms*.

## Notes

### Competing Interest Statement

The authors have declared no competing interest.

