## Supplementary figures and images for "Encoding social preference by interhemispheric neurons in the Insula"

### supplementary figures

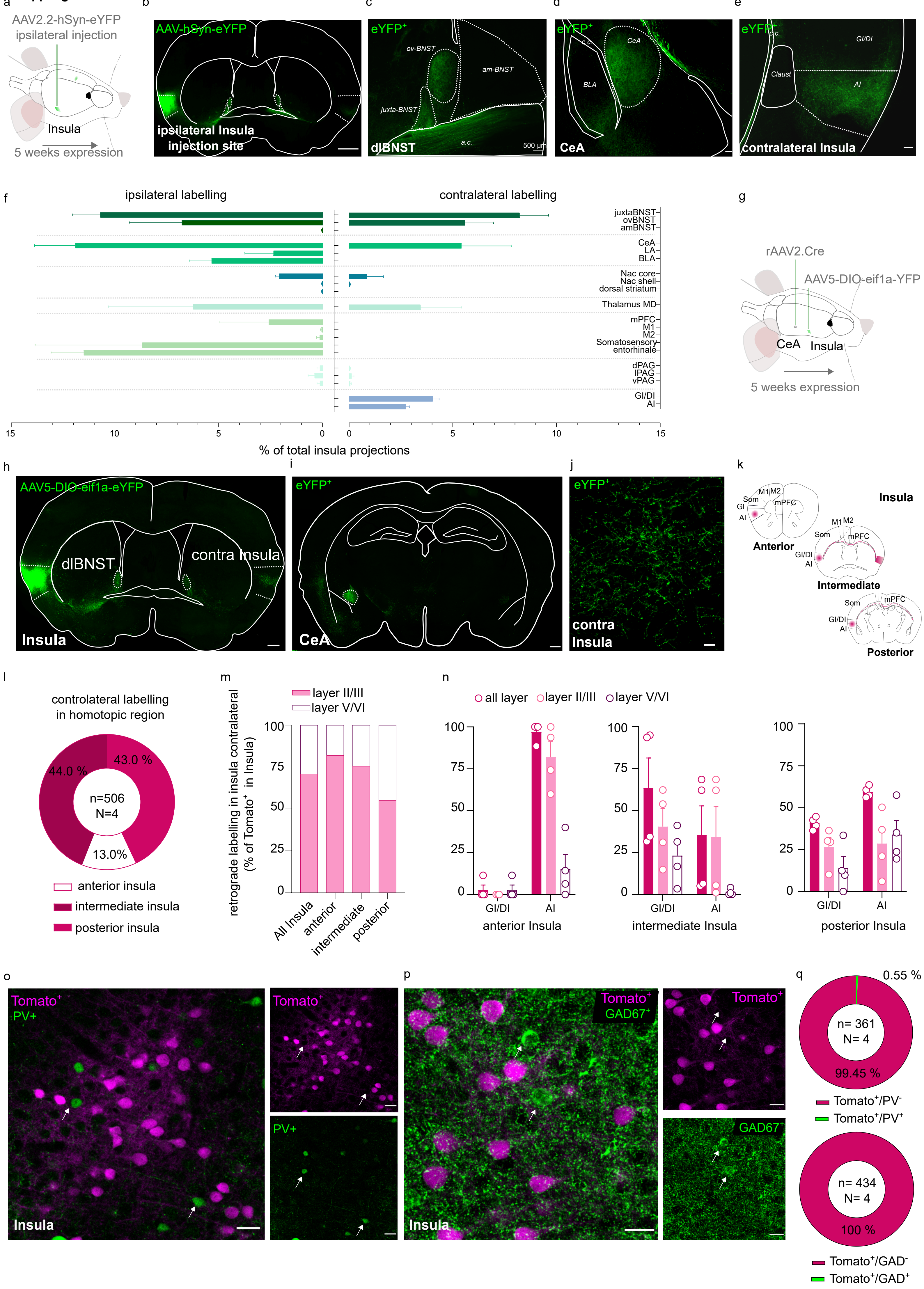

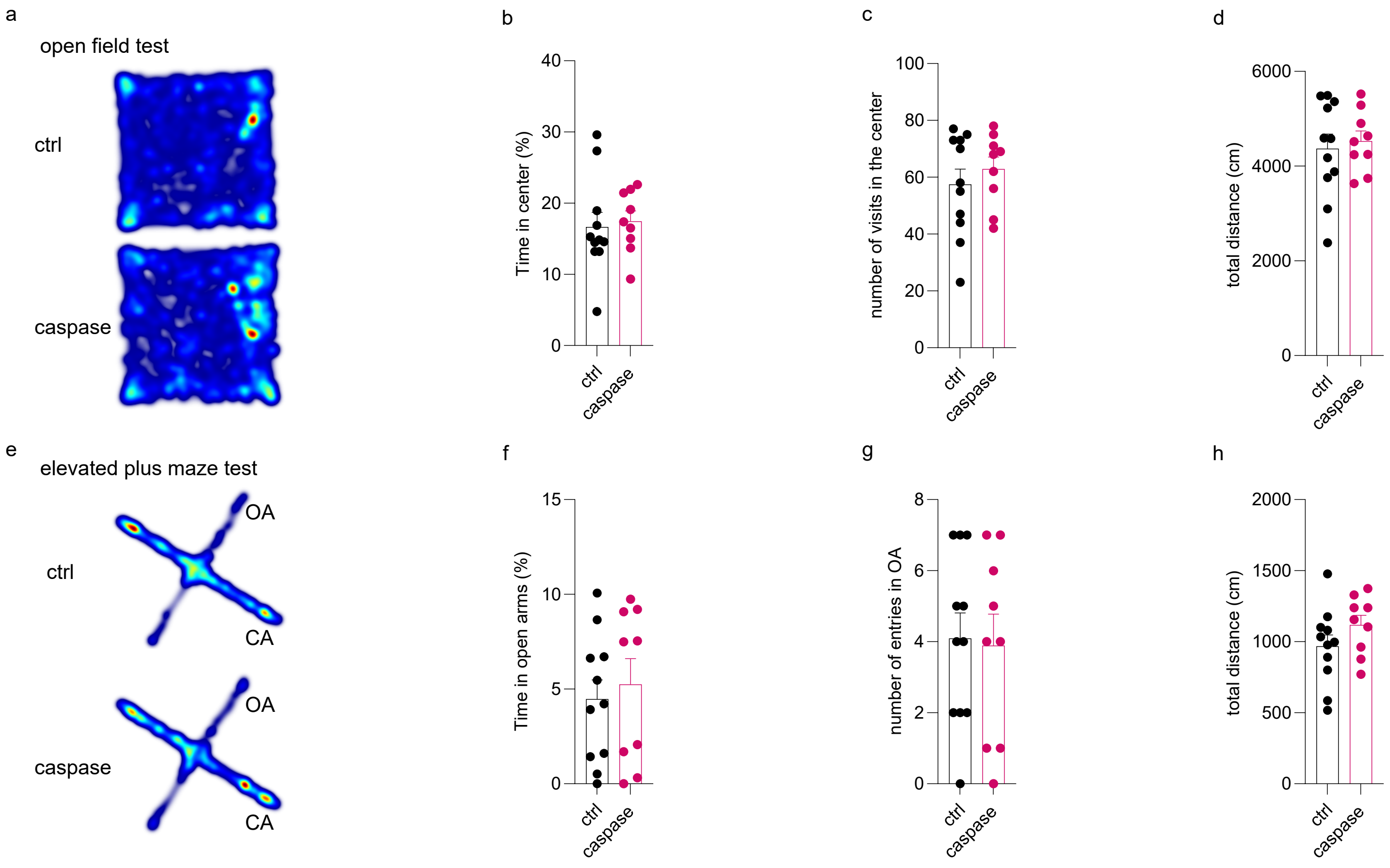
